# Time-course Transcriptomics Reveals the Impact of *Treponema pallidum* on Microvascular Endothelial Cell Function and Phenotype

**DOI:** 10.1101/2025.05.04.651946

**Authors:** Sean Waugh, Mara C. Goodyear, Alloysius Gomez, Akash Ranasinghe, Karen V. Lithgow, Reza Falsafi, Robert E.W. Hancock, Amy H. Lee, Caroline E. Cameron

## Abstract

Syphilis, caused by *Treponema pallidum* subsp. *pallidum,* is an urgent global public health threat. Syphilis vaccine development has been impeded by limited understanding of the molecular mechanisms that enable *T. pallidum* to establish and maintain infection. The vascular endothelium is critical for *T. pallidum* attachment, dissemination, and host immune response initiation; however, the molecular details of *T. pallidum*-endothelial interactions are incompletely understood. To enhance understanding, we performed time-course transcriptomic profiling on *T. pallidum*-exposed brain microvascular endothelial cells. These analyses showed *T. pallidum* exposure altered pathways related to extracellular matrix, growth factors, integrins, and Rho GTPases. The induced transcriptional response was consistent with endothelial to mesenchymal transition, a process involved in fetal development and vascular dysfunction. This study provides a comprehensive understanding of the molecular responses of endothelial cells to *T. pallidum* and identified the host pathways that might cause syphilis disease symptoms, information that could aid in syphilis vaccine design.

**Highlights:**

□ Exposure of microvascular endothelial cells to *Treponema pallidum* subsp. *Pallidum* significantly alters the endothelial cell transcriptome
□ Signaling pathways related to extracellular matrix organization, growth factors, integrins, and Rho GTPases were overrepresented for genes differentially expressed in *T. pallidum*-exposed endothelial cells
□ Exposure to *T. pallidum* induces pathways and factors consistent with endothelial to mesenchymal transition, a host process central to development that may explain the devastating effects of congenital infection
□ *T. pallidum* exposure induces expression of Snail, the main transcription factor associated with the process of endothelial to mesenchymal transition

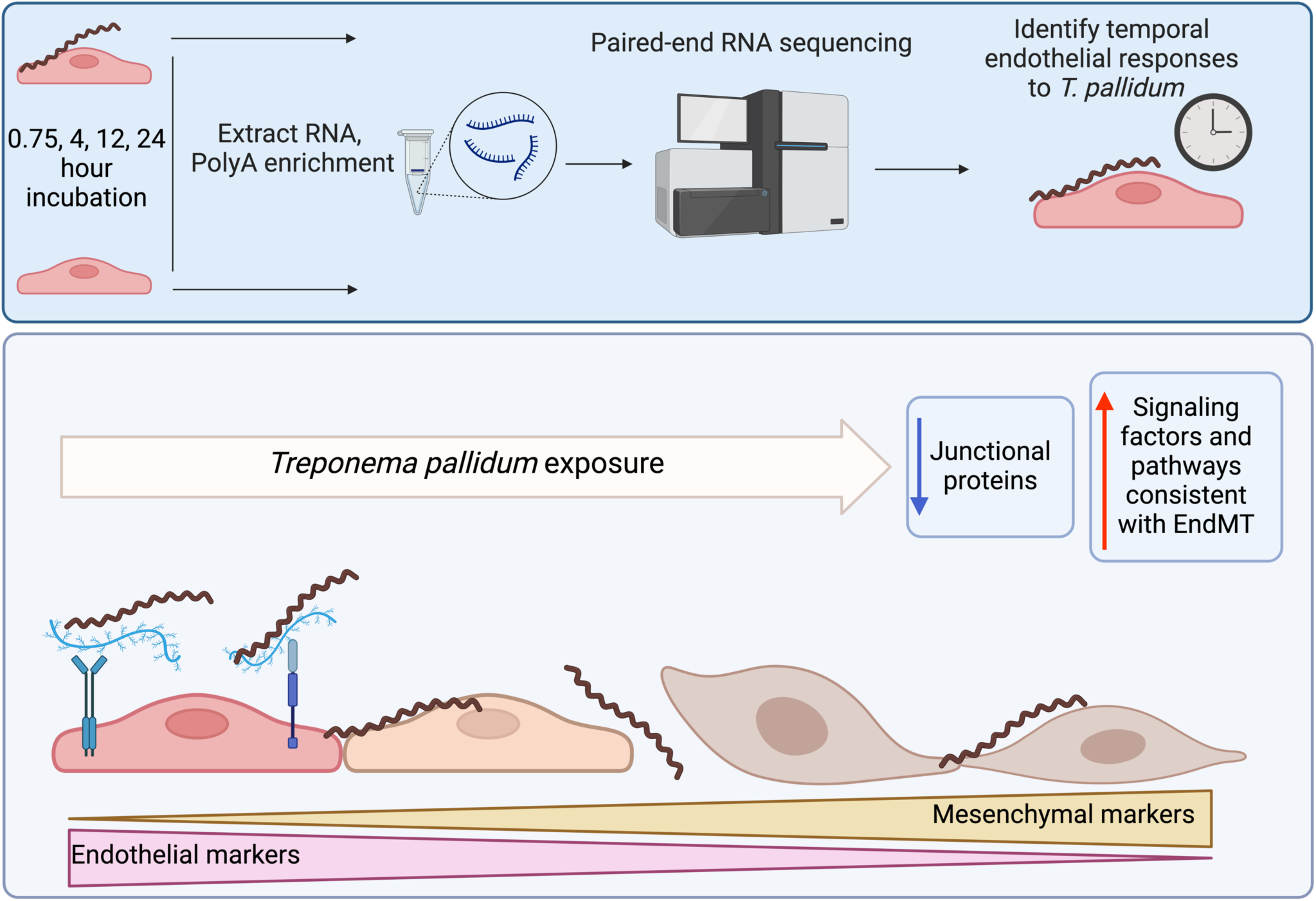

## Introduction

Infectious syphilis, caused by the sexually transmitted bacterium *Treponema pallidum* subsp. *pallidum*, is a multi-stage infection with a global burden of 49.7 million cases^1^ and 8 million new infections per year amongst individuals 15-49 years of age^2^. The infection is systemic, and persists for an individual’s lifetime in the absence of effective antibiotic treatment^3^. The bacterium also causes congenital syphilis, with the number of congenital syphilis cases worldwide estimated at approximately 661,000 resulting in 355,000 adverse birth outcomes per year^4^. These figures likely represent an underestimation since an accurate determination of the burden of congenital syphilis worldwide is challenging, due to variations in antenatal screening coverage, access to syphilis testing during pregnancy, and availability and quality of surveillance data. As global syphilis cases have reached a 20-year high^1,5–7^, understanding the molecular basis of the *T. pallidum*-host interaction is crucial for developing biomedical interventions to address the escalating public health burden posed by this pathogen.

*Treponema pallidum* is a highly invasive pathogen with the ability to disseminate via the bloodstream and cross endothelial, blood-brain, and placental barriers^3^. Interactions between *T. pallidum* and microvascular endothelial cells are central to both *T. pallidum* dissemination via the bloodstream and the establishment of disease manifestations. Previous studies investigating endothelial responses to *T. pallidum* using targeted molecular approaches have provided insights into the cellular consequences of *T. pallidum*-host interactions^8–12^. Global proteomics and immune secretomics analyses of *T. pallidum-* exposed human brain microvascular endothelial cells (HBMECs) revealed numerous changes in endothelial cellular signaling pathways, including pathways involved in extracellular matrix (ECM) structural changes, dysregulation of cell death/necroptosis, induction of pro-inflammatory cytokine profiles, and suppression of macrophage/monocyte activating immune responses^13^.

To expand on these findings and identify host signaling pathways that play a role in syphilis pathogenesis, the current study characterized HBMEC transcriptional responses to *T. pallidum* exposure by time-course transcriptome sequencing (RNA-seq) conducted at four timepoints over a sampling period of 45 minutes to 24 hours post-exposure. These analyses reveal significant pathway convergence on endothelial to mesenchymal transition (EndMT), a dynamic process involved in various developmental, physiological, and pathological activities^14,15^ that contributes to vascular destruction in other infectious diseases, such as SARS-CoV-2 infection^16^. These characteristics of EndMT suggest potential relevance to the development of symptoms associated with infectious and congenital syphilis. In the current study, differential expression of growth factor signaling pathways and genes related to the process of EndMT were enriched in the obtained dataset. This includes the upregulation of *SNAI1* (Snail), a transcription factor which serves as the central transcriptional regulator of EndMT progression^17^ and has been shown to contribute to blood-brain-barrier (BBB) disruption during pathogen contact^18,19^. The systems biology approach pursued in this study provides insight into molecular signaling and cellular transformations occurring in the host that collectively shape the course of *T. pallidum* infection.

## Results

### Endothelial transcription was significantly altered during *T. pallidum* exposure

To identify HBMEC genes that were differentially expressed (DE) during *T. pallidum* exposure, RNA-Seq was conducted on five independent wells per treatment condition comparing a viable *T. pallidum* (VTP) exposure to an infection extract control (IEC) exposure at each timepoint. Briefly, these conditions corresponded to exposure to a VTP sample containing viable *T. pallidum,* alongside exposure to an IEC sample where *T. pallidum* was removed using a centrifugation and filtration technique that retains background culture contaminants co-purified during *T. pallidum* harvest^13^. Exposing HBMECs to *T. pallidum* significantly altered gene expression in the brain endothelial cells, where the number of DE genes with adjusted p-value of < 0.05 and a fold-change (FC) cutoff of ±1.5 were 11 at 45-minutes, 225 at 4-hours, 490 at 12-hours, and 1753 at 24-hours (Figure 1A-D; Supplementary Table 1). Considerable overlap in DE genes was observed between timepoints, where 311 genes were differentially expressed at both 12 and 24 hours, 52 genes were differentially expressed from 4-24 hours, and 4 genes were differentially expressed at all timepoints (Figure 1E).

**Figure 1.**
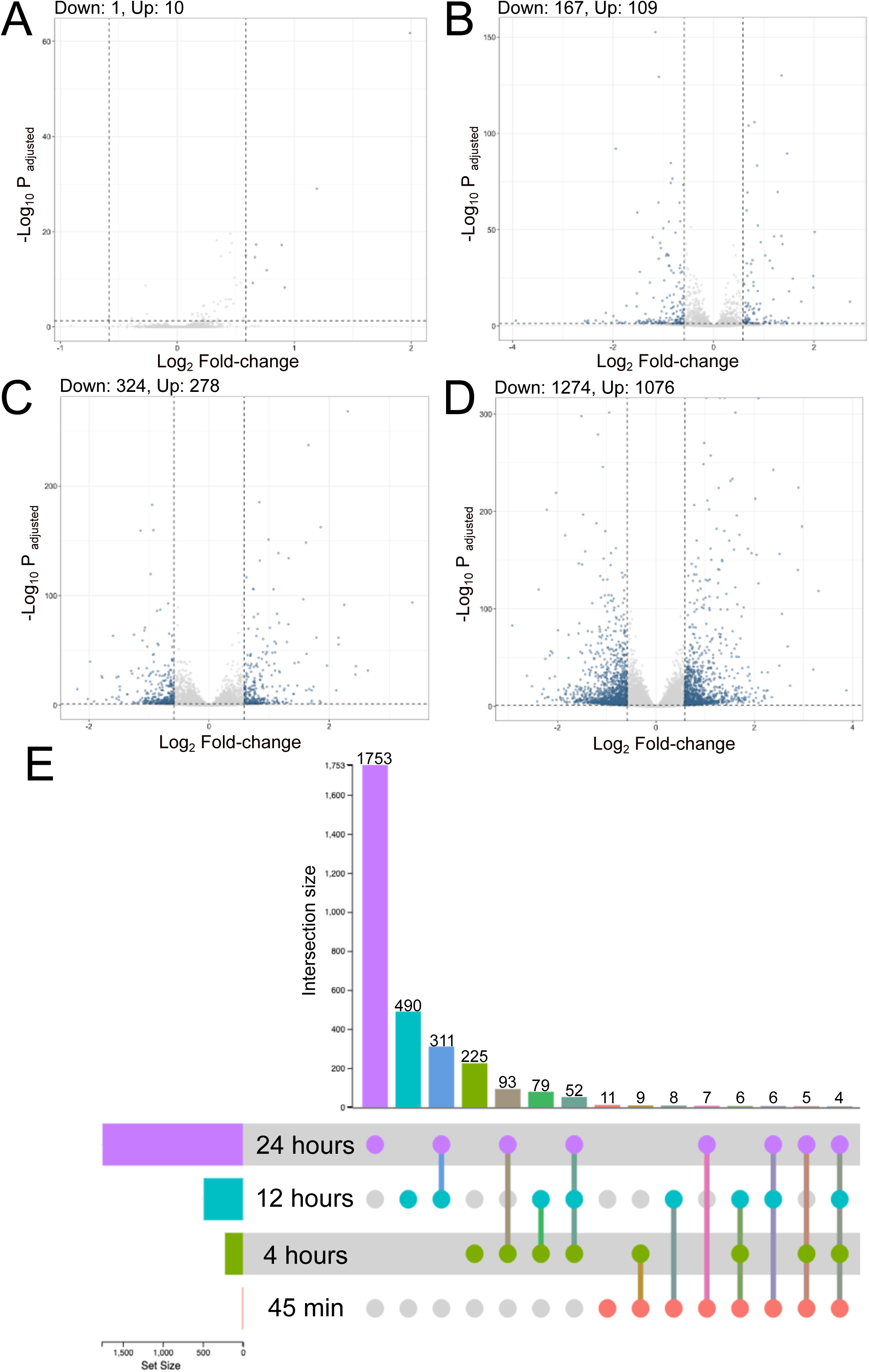
The number of DE genes in VTP versus EC-exposed cells increased with longer *T. pallidum* exposure times, and DE genes demonstrated significant overlap between timepoints. Volcano plots showing DE genes at (A) 45-minute, (B) 4-hour, (C) 12-hour, and (D) 24-hour exposures. Dotted lines denote a fold-change cutoff of ±1.5, and an adjusted p-value cutoff of 0.05. Significant DE genes are dark grey, non-significant genes are light grey. The number of up- and downregulated genes per timepoint is shown above the plot. (E) Upset plot showing the intersection of DE genes between timepoints.

### *Treponema pallidum* exposure altered expression of transcription factors associated with cellular differentiation

Next we completed Transcription factor (TF) enrichment and co-regulatory analyses^20^ on each of the study timepoints to identify TFs that may be responsible for the observed changes in gene expression in HBMECs during *T. pallidum* exposure. Using an integrated mean ranking algorithm^20^, local TF co-regulatory networks were generated for the top 10 TFs predicted to be most influential in regulating the expression of the DE genes at each timepoint (Figure 2A-D; Supplementary Tables 2-5). These analyses identified highly connected co-regulatory TF networks at each timepoint, all of which featured AP-1 (Activating Protein-1) subunits such as Fos, FosB, and Jun (Figure 2A-D). Notably, the transcripts for *FOS*, *FOSB*, *JUN*, *JUNB*, and *JUND* were upregulated at one or more timepoints in this study (Supplementary Table 1). NR4A1 (Nuclear Receptor subfamily 4 group A member 1) and NR4A3 were also enriched at the 12- and 24-hour timepoints, and transcripts encoding these TFs were significantly upregulated at all timepoints in the study. Transcription factors involved in the cellular differentiation processes of EndMT and Epithelial to mesenchymal transition (EMT) were also enriched at all timepoints, including HEY1 (Hes-related family bHLH transcription factor with YRPW motif 1), HEY2, BHLHE40 (Basic Helix-Loop-Helix Family Member e40), EGR1 (Early Growth Response 1), and EGR2. Snail, the primary TF driving EndMT^21–23^ ranked among the top 10 most enriched TFs for all timepoints (Supplementary Tables 2-5). Significantly increased *SNAI1* expression was observed in VTP-exposed HBMECs from 12- to 24-hours (3.08, 3.11 FC, respectively), contrasting with minimal alteration of *SNAI1* expression in IEC-exposed HBMECs over the same period (Figure 3A). Western blot validation confirmed *T. pallidum* exposure induced Snail protein upregulation, with significantly higher protein abundance at 24 hours and increased protein abundance at 12 hours (VTP-versus IEC-exposed cells; Figure 3B-C). Collectively, these findings identify the early and sustained alteration of EndMT-associated TFs and their influence on HBMEC transcriptional responses to *T. pallidum* exposure.

**Figure 2.**
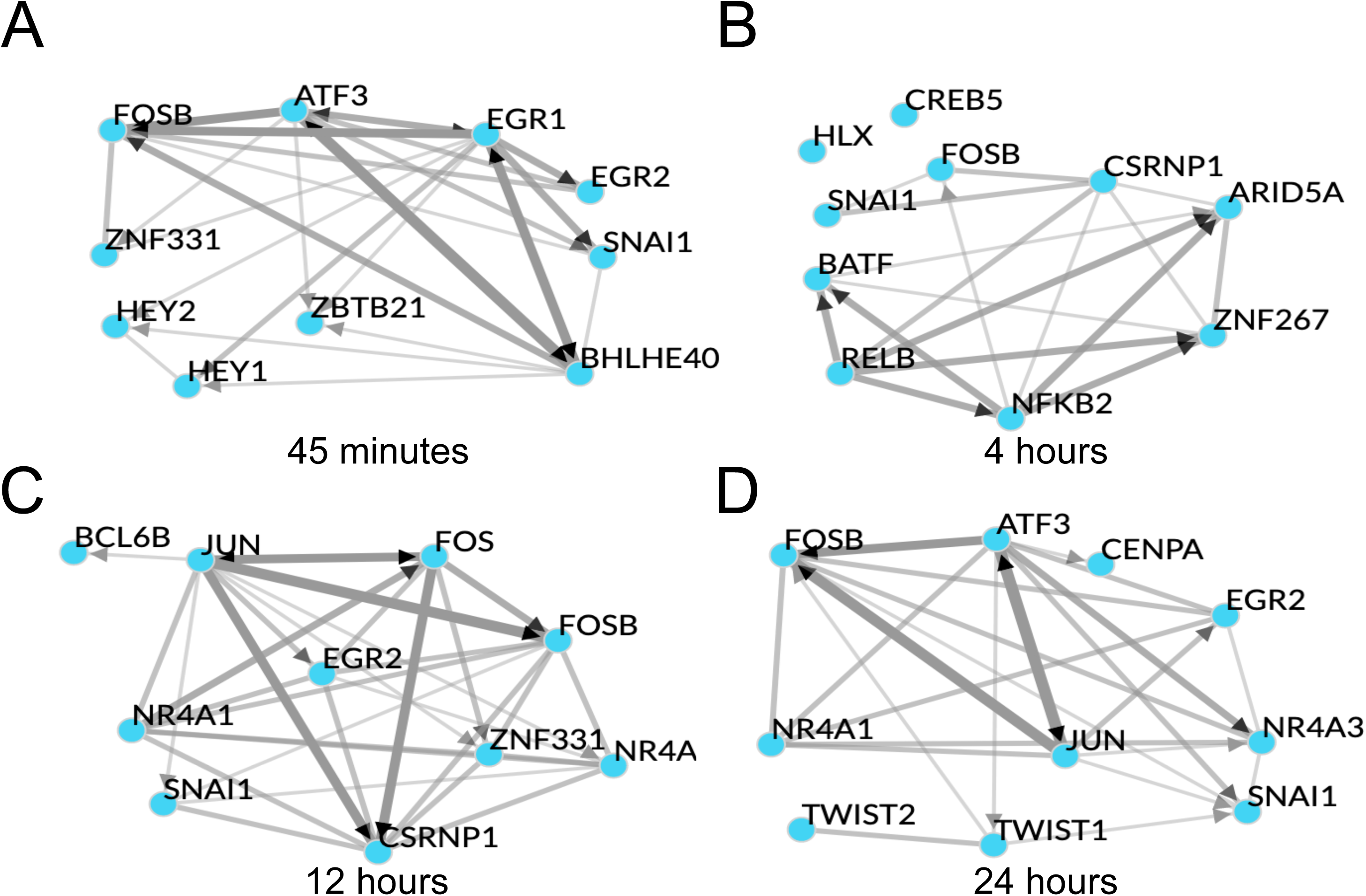
Transcription factor enrichment revealed the top 10 most enriched TFs at each timepoint form highly connected co-regulatory networks. Shown are Chea3 transcription factor enrichment integrated mean analyses for the timepoints (A) 45-minutes, (B) 4-hours, (C) 12-hours, (D) 24-hours. Edge (line) width represents the strength of interaction from chromatin immunoprecipitation sequencing (ChIPseq), co-expression, and co-occurrence data. Black arrowheads indicate the direction of co-regulation where ChIP-seq supports the direction of regulation, and undirected (no arrows) where only co-occurrence or co-expression data is available.

**Figure 3.**
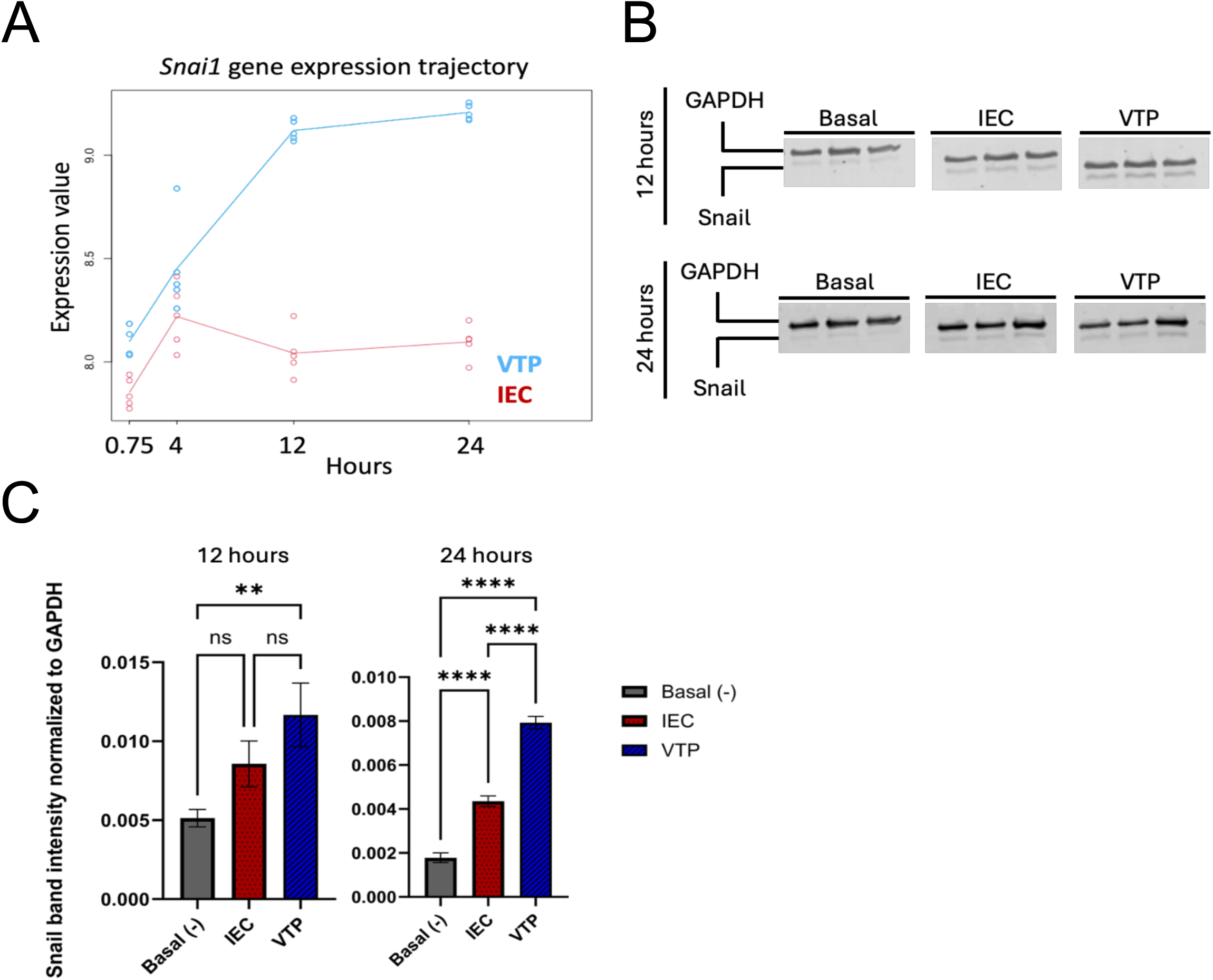
*SNAI1* gene and Snail protein expression was increased in HBMECs exposed to *T. pallidum*. (A) *SNAI1* (Snail) longitudinal expression plot showing Log-transformed expression values at each timepoint for viable *T. pallidum* exposed cells (VTP; Blue) versus infection extract control (IEC; Red). Each data point represents an individual biological replicate. Analysis and figure generation completed using maSigPro (v1.74.0). (B) Temporal analysis of Snail protein expression in endothelial cells exposed to VTP, IEC, and Basal Media conditions. Each condition comprises three biological replicates, defined as individual tissue culture wells exposed to the treatment conditions. GAPDH was used as a loading control and for normalization. The 12- and 24-hour timepoints were run on separate gels. Empty lanes separating different sample conditions (Basal, IEC, VTP) were removed. (C) Quantification of Snail protein abundance normalized to GAPDH as a loading control. Quantitation was completed on a Licor Odyssey CLx using Licor Image Studio version 5.2. Significant differences in Snail abundance determined by One-way ANOVA followed by Tukey’s multiple comparisons test, where comparisons were considered significant with a p-value ≤ 0.05. * indicates a P-value ≤ 0.05, ** indicates a P-value ≤ 0.01, *** indicates a P-value ≤ 0.001, and **** indicates a P-value ≤ 0.0001.

### *Treponema pallidum* exposure altered expression of pathways within the categories of gene expression, extracellular matrix organization, immune system, programmed cell death, and signal transduction

We also performed Sigora pathway overrepresentation analysis^24^ on up- and down-regulated genes at each timepoint to investigate pathways that are significantly overrepresented in *T. pallidum*-exposed HBMECs. Sigora uses Reactome database annotations^25^ and contextualizes pathway overrepresentation by weighting DE gene pairs that occur uniquely in a single pathway. The number of overrepresented pathways increased with time, with 1, 15, 19, and 46 significantly overrepresented pathways at 45-minutes, 4-, 12-, and 24-hours respectively. These investigations identified overrepresented cellular pathways within the categories of gene expression, extracellular matrix (ECM) organization, immune system, programmed cell death, disease, and signal transduction, and key pathways from these categories are highlighted below (Figure 4A-D; Supplementary Tables 6-9).

**Figure 4.**
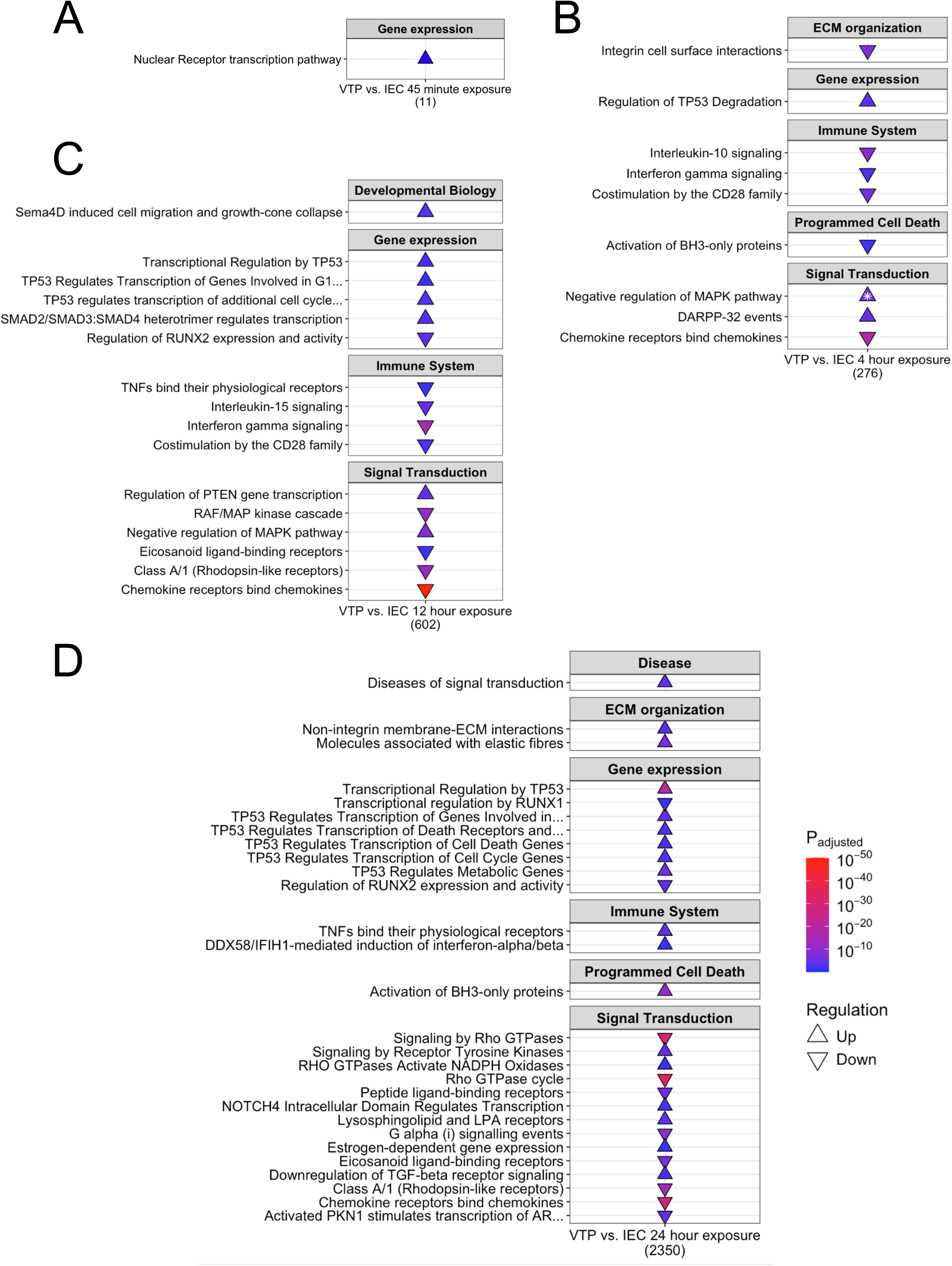
*Treponema pallidum* exposure altered expression of genes involved in the categories of gene expression, extracellular matrix organization, immune system, programmed cell death, and signal transduction. Shown is a subset of overrepresented pathways in HBMECs exposed to *T. pallidum* that were upregulated (Δ) and downregulated (∇) at the timepoints of (A) 45-minutes, (B) 4-hours, (C) 12-hours, (D) 24-hours. The total number of DE genes for each timepoint is listed under each table. White stars on the pathway indicate that the pathway was both up- and downregulated, with the direction of arrow indicating the most statistically significant direction of regulation. The subset of overrepresented pathways included align with overarching functional categories described in the results. The full list of ovverepresented pathways are listed in Supplementary Tables 6-9.

Within the category of gene expression, we observed overrepresentation of pathways related to p53-mediated cell death and p53-mediated transcriptional regulation at the 4-to 24-hours timepoints (Figure 4B-D; Supplementary Tables 7-9). At the 12-hour timepoint, pathways related to EndMT were upregulated, such as “SMAD2/SMAD3:SMAD4 transcriptional regulation” and “regulation of PTEN gene expression” (Figure 4B; Supplementary Table 7). We also observed dysregulation of pathways involved in ECM organization, with “integrin signaling” first downregulated at 4-hours, followed by upregulation of additional ECM pathways (“non-integrin membrane-ECM interactions”, “molecules associated with elastic fibres”, and “signaling by receptor tyrosine kinases”) at 24-hours (Figure 4B, 4D; Supplementary Tables 7, 9). Within the category of immunity, we observed downregulation of pathways in “chemokine receptors bind chemokines” and TNF and IFN signaling (4- and 12-hours), suggesting inflammatory pathway downregulation (Figures 4B, 4C). While “chemokine receptor signaling” remained downregulated at 24-hours, pathways involving TNF and IFN signaling were upregulated, indicating differential temporal regulation of these immune pathways in HBMECs exposed to *T. pallidum* (Figure 4D). Similarly, the cell death signaling pathway “activation of BCL2-Homology 3 (BH3)-only proteins” was downregulated at 4-hours but upregulated at 24-hours (Figure 4B, D). Within the category of signal transduction, we observed the pathway “negative regulation of MAPK (Mitogen-Activated Protein Kinase)” to be upregulated at 4- and 12-hours, and “RAF/MAP kinase cascade” to be downregulated at 12-hours, indicating the involvement of MAPK signaling in the endothelial response to *T. pallidum.* At the 24-hour timepoint, overrepresented pathways within the category of signal transduction included “downregulation of TGFβ receptor signaling”, and upregulation of “NOTCH4 intracellular domain regulates transcription”, “signaling by receptor tyrosine kinases”, and “diseases of signal transduction” (Figure 4D). Pathways related to Rho GTPase activity were downregulated at 4- and 24-hours, however the downregulated genes in these pathways are primarily GTPase-activating proteins (GAPs) which negatively regulate Rho GTPases (Supplementary Tables 7, 9). Further, the Rho GTPase-mediated pathway “Sema4D induced cell migration and growth cone collapse” and “Rho GTPase activates NADPH oxidases” were upregulated at 12- and 24-hours, respectively (Figure 4C, D). To identify temporal connections between overrepresented pathways, we next performed pathway network analysis at each timepoint, to identify pathway clusters and connections based on overlapping DE genes (Figure 5A-D). Through this analysis we determined that the relationships between overrepresented pathways became more connected with longer *T. pallidum* exposures (Figure 5A-D). At earlier timepoints, pathways formed distinct clusters which were separated into categories such as immune responses and signal transduction (Figure 5A-C). The 24-hour network showed the most connections within and between clusters of pathways, including connections between clusters belonging to different categories (Figure 5D), suggesting a converging transcriptional trajectory in *T. pallidum*-exposed HBMECs.

**Figure 5.**
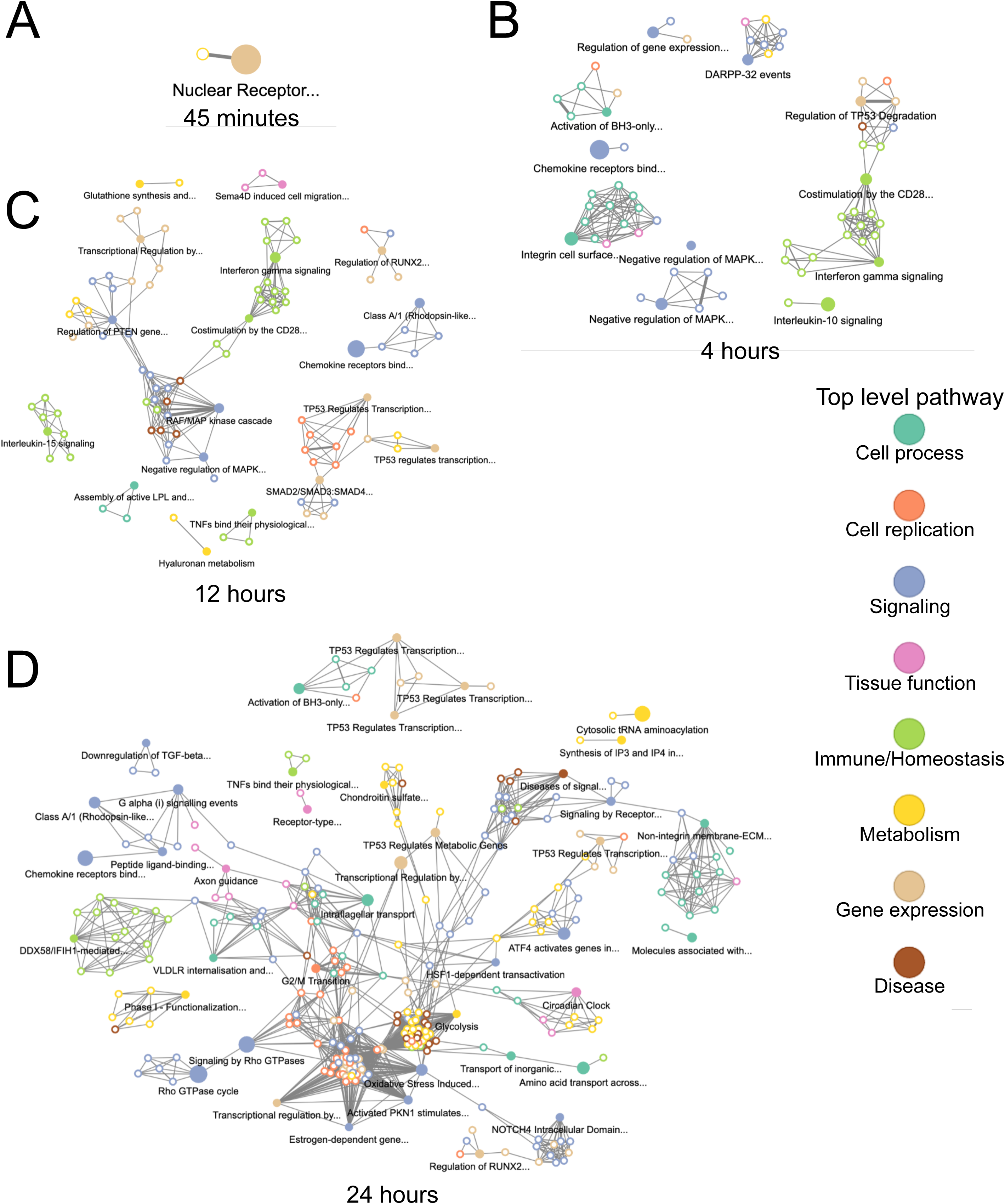
Overrepresented pathways in *T. pallidum*-exposed HBMECs became more connected with increased exposure duration. Visualization of networks of overrepresented Reactome pathways at (A) 45-minutes, (B) 4-hours, (C) 12-hours, (D) 24-hours of exposure. Pathway connection is determined using the overlap of the genes assigned to each pathway to determine their similarity to one other. Each node represents a pathway, and the connection/edge connecting nodes was determined using a maximum Jaccard distance of 0.85. Significantly overrepresented pathways are represented by a solid-colored node and have an associated pathway title. The nodes with a colored border and white core are not significantly overrepresented. Nodes are colour-coded according to the legend by top-level pathway. The connections between pathways are represented by the thickness of the edge connecting each node.

### *Treponema pallidum* exposure resulted in overrepresentation of hallmark gene sets in the categories of development, immune responses, cellular signaling, and cellular stress

To identify higher-order overrepresented transcriptional profiles in HBMECs exposed to *T. pallidum,* we performed hallmark molecular signature enrichment analysis using the MSigDB database^26^ on each study timepoint. Hallmark gene sets are curated and contextual, whereby gene function is considered for direction of regulation. Due to this context, and the higher-order approach to pathway analysis, hallmark gene sets can be both up- and downregulated at single timepoints. Through these analyses, we identified hallmark gene set overrepresentation within the categories of development, immune, signaling, and stress (Figure 6A-D), supporting the results of the Sigora pathway overrepresentation analysis. Consistent with the pathway analysis, hallmark gene set enrichment also identified inflammatory gene sets to be downregulated. Additionally, the gene set “TNF-α signaling through NF-κB” was simultaneously up- and downregulated at all timepoints, resulting from concurrent differential regulation of genes belonging to these pathways (Figure 6A-D). Further, the “apoptosis” gene set was overrepresented at all timepoints, although the direction of regulation varied (Figure 6B-D). We also observed overrepresented gene sets within the signaling category, including downregulation of “KRAS signaling UP” from 4-to 24-hours, and upregulation of both “Wnt β-catenin signaling” and “NOTCH signaling” at 24-hours (Figure 6B-D).

**Figure 6.**
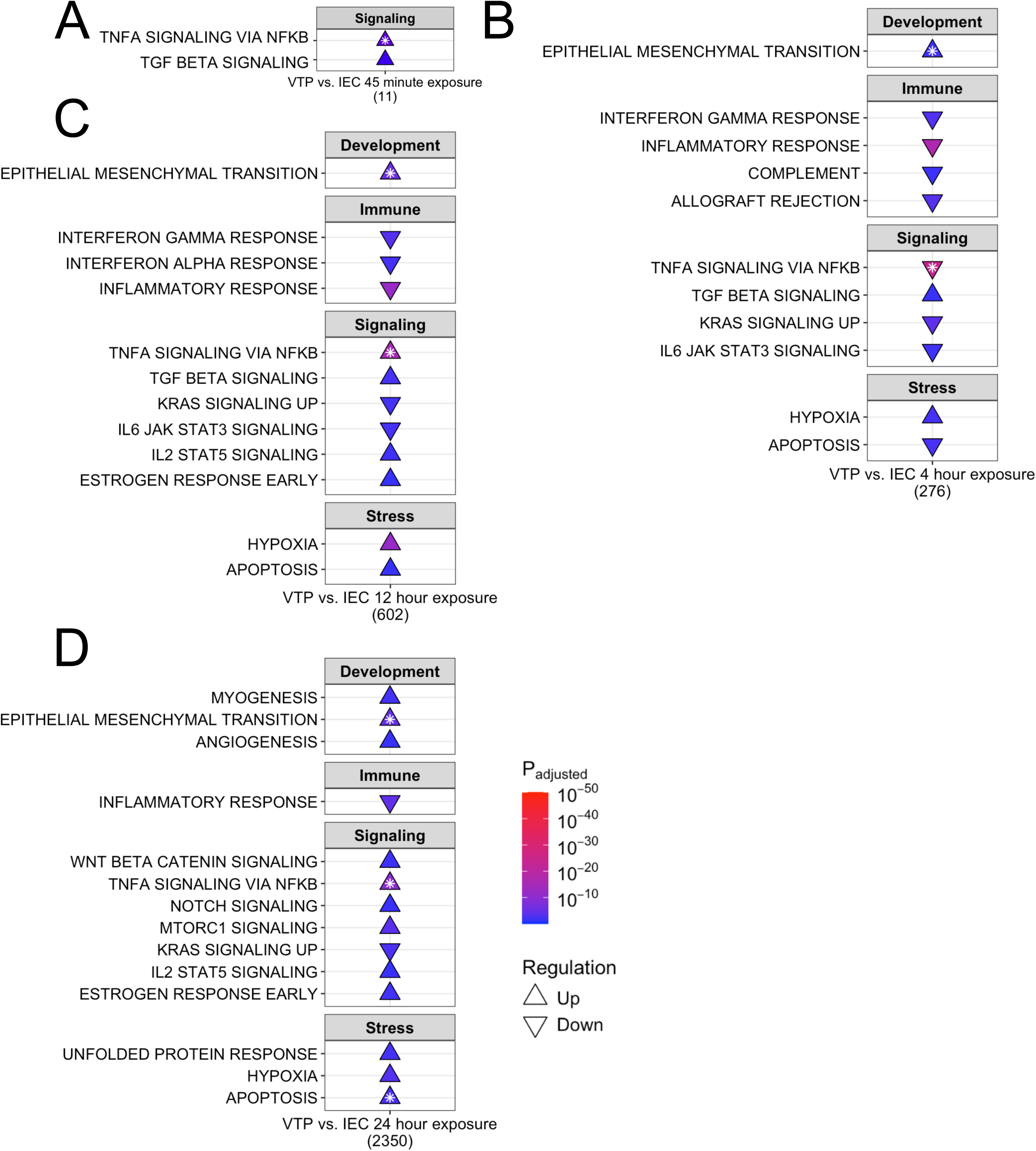
Hallmark gene categories of development, immune responses, cellular signaling, and stress were enriched in *T. pallidum*-exposed HBMECs. Shown is a subset of overrepresented hallmark gene sets in HBMECs exposed to *T. pallidum* that were upregulated (Δ) and downregulated (∇) at the timepoints of (A) 45-minutes, (B) 4-hours, (C) 12-hours, (D) 24-hours. The total number of DE genes for each timepoint is listed under each table. White stars on the pathway indicate that the pathway was both up- and downregulated, with the direction of arrow indicating the direction of regulation with the lowest adjusted p-value. The subset of gene sets included align with overarching functional categories described in the results. The full list of overrepresented hallmark genesets are listed in Supplementary Tables 10-13.

In support of the overrepresented TGFβ and SMAD signaling pathways identified by Sigora pathway analysis, in the hallmark gene set “TGFβ signaling” was also upregulated from 45-minutes to 12-hours (Figure 6A-C). Notably, the “epithelial to mesenchymal transition (EMT)” gene set was both up- and downregulated from 4- to 24-hours, with upregulation being most significant at these timepoints (Figure 6A-D). This bi-directional regulation reflects the dynamic regulation of this cellular process. Currently, there is no hallmark gene set for EndMT; however, EMT and EndMT have shared regulatory pathways^27^, and thus this analysis provides insight into the regulatory factors common to both pathways. Additional EMT- and EndMT-related hallmark gene sets were overrepresented at the 24-hour timepoint, such as “myogenesis” and “angiogenesis” (Figure 6D; Supplementary Table 13). These data reinforce that EndMT-inducing pathways and gene sets, such as TGFβ, SMAD, and NOTCH signaling, and *SNAI1* transcription/Snail protein expression were induced in HBMECs exposed to *T. pallidum*.

We observed that many genes involved in EndMT-related processes were DE, including upregulation of the SMAD negative regulator *PMEPA1* (Prostate transmembrane Protein Androgen Induced 1) and inhibitory *SMAD7*, as well as downregulation of activating *SMAD3*. Genes associated with *SNAI1* overexpression^28^ were also DE, such as upregulation of the EndMT-promoting ECM component *FBLN5* (Fibulin-5), and downregulation of adhesion molecules^29^ *ICAM-1* (Intercellular Adhesion Molecule 1), *VCAM-1* (Vascular Cell Adhesion Protein 1), as well as genes encoding junctional proteins, including *CLDN1* (Claudin-1), *OCLN* (Occludin), and *ZO-2* (Zona Occludens-2 [also known as *TJP2*]) (Supplementary Table 1). *vWF* (von Willebrand Factor), a secreted factor important for endothelial inflammation, angiogenesis, endothelial dysfunction, and adhesion of platelets and leukocytes to endothelial cells^30^, was upregulated at the 12- and 24-hour timepoints (Supplementary Table 1). Mesenchymal marker expression^14^ was increased, including upregulation of *SM22-α* (Smooth Muscle Protein 22 Alpha [also called *TAGLN*]), *α-SMA* (Smooth Muscle Actin Alpha 2 [also called *ACTA2*]), *Integrin-β3*, *ACTC1* (Actin Alpha Cardiac Muscle 1), *COL3A1* (Collagen type III A1), *COL4A1*, *COL4A2*, *POSTN* (Periostin), and *FN1* (Fibronectin) (Supplementary Table 1). These data demonstrate that HBMECs exposed to *T. pallidum* undergo significant transcriptional alterations consistent with the process of EndMT, and identified the involvement of Snail in the host endothelial response to *T. pallidum*.

### Growth factor pathways and genes involved in EndMT are altered in HBMECs exposed to *T. pallidum*

We found that HBMECs exposed to *T. pallidum* demonstrated transcriptional alteration indicative of TGFβ and SMAD signaling, as well as pathways that modify TGFβ signaling such as NOTCH1, Wnt/β-catenin, and receptor tyrosine kinase (RTK) signaling, all of which contribute to EndMT (Figures 4, 6; Supplementary Tables 6-13). Importantly, TGFβ signaling is controlled through feedback loops, particularly negative feedback, where genes induced by TGFβ signal activation both attenuate the activity of TGFβ signaling proteins and repress transcription of genes involved in TGFβ signaling^31^. The findings in the current study are consistent with the activation and downstream regulation of TGFβ signaling. Specifically, Sigora pathway analysis identified upregulation of the pathway “SMAD2/SMAD3:SMAD4 heterotrimer regulates transcription” at 12-hours, which is the primary transcriptional response induced by TGFβ signal activation^31^ (Figure 5C). Negative feedback was also evident through upregulation of the pathway “downregulation of TGFβ receptor signaling” at 24-hours, where the DE genes in this pathway are upregulated as a negative feedback response to TGFβ signal activation (Figure 6D). Further, hallmark gene set analysis, which accounts for the context and function of DE genes, identified upregulation of TGFβ signaling from 45-minutes to 12-hours (Figure 6A-C). These data indicate the activation and regulation of TGFβ signaling pathways in the endothelial response to *T. pallidum*.

To further investigate these findings, we assessed the expression changes of specific growth factor genes involved in EndMT-activating and regulatory pathways. Specific DE genes within TGFβ signaling include the downregulation of *TGFBR* (TGFβ Receptor)-*2* and -*3* at 12- and 24-hours, and *SMAD3*, a prominent activating TF downstream of TGFβ signaling, at the 4- and 12-hour timepoints (Supplementary Table 1). Expression of TGFBRs is negatively regulated by genes induced by TGFβ signaling, such as the inhibitory *SMAD*7, which is induced by the SMAD2/SMAD3:SMAD4 dimer as a negative feedback mechanism^31,32^ and was upregulated in this study (Supplementary Table 1). Upregulation of additional TGFβ-induced negative regulators^31^ was observed, including *BAMBI* (BMP and Activin Membrane Bound Inhibitor) at 24-hours, and *PMEPA1* at 12- and 24-hours (Supplementary Table 1). Additional upregulated TGFβ-induced genes^22,33^ include *ID* (Inhibitor of DNA binding)*-1*, -*2*, and -*3*, and fibrosis regulator^34^ *CCN2* (Cellular Communication Network Factor 2) (Supplementary Table 1). Upregulated genes that enhance TGFβ and EndMT include *EDN1* (Endothelin-1) at 4- to 12-hours, and *POSTN* (Periostin) that is involved in TGFβ-induced EndMT/EMT through integrin-β3 signaling^35^, which was upregulated at 24-hours. These data demonstrate the activation and regulation of TGFβ signaling in HBMECs exposed to *T. pallidum*.

TGFβ can also signal independently of SMADs, including through pathways involving MAPK cascades, GTPases, and Wnt/β-catenin^14^. In the current study, we observed significant overrepresentation of these signaling pathways at one or more timepoints, as determined by both Sigora and hallmark gene set overrepresentation analyses (Figures 4, 6; Supplementary Tables 6-13). Among these pathways, *DUSP2* (Dual Specificity Phosphatase 2) and *DUSP8* were among the most upregulated genes, indicating negative feedback of MAPK signaling. Other regulators in the EndMT and non-canonical TGFβ pathways include the upregulation of *NECTIN4* and *IL-11*, the latter of which contributes to fibrosis and EndMT and was upregulated at all timepoints^36,37^. Correspondingly, *NOX4* (NADPH oxidase 4), which mediates fibrosis and cellular differentiation downstream of IL-11 and TGFβ^38^, was upregulated from 4- to 12-hours. Additionally, *NEDD9* (neural precursor cell expressed, developmentally down-regulated 9), a focal adhesion scaffold protein involved in transducing cell adhesion and integrin signaling through RTKs, as well as promoting EMT and cancer metastasis^39^, was significantly upregulated at 12 and 24 hours. Other growth factor signaling genes that contribute to EndMT were also upregulated, including *PDGFB* (Platelet-Derived Growth Factor B), *VEGFA* (Vascular Endothelial Growth Factor A), *PGF* (Placental Growth Factor), and *FLT1*, which encodes VEGF Receptor 1 (VEGFR1). In alignment with the differential expression of growth factors and overrepresentation of growth factor signaling pathways and hallmark gene sets, RTK signaling pathways were significantly overrepresented at the 24-hour timepoint (Figure 4D). Collectively, these data

demonstrate the alteration and regulation of growth-factor signaling genes and pathways in HBMECs during *T. pallidum* contact.

### ECM regulatory factors, components, and pathways were dysregulated during *T. pallidum* exposure

In the current study, we found *T. pallidum*-exposed HBMECs exhibited significant transcriptional alterations of ECM constituents and regulatory proteins, accompanied by overrepresentation of ECM organization and signaling pathways at several timepoints (Figure 4B, D; Supplementary Tables 7, 9). ECM components that were DE included upregulation of collagen subtypes *3*, *4*, *5*, *7* and *27*, and downregulation of *COL5A3* (Supplementary Table 1). The collagen receptor and EMT positive regulator^40^, *DDR2* (Discoidin Domain Receptor 2), was also upregulated at 24-hours, as were the ECM components *LAMA1* (Laminin subunit alpha 1) and *FN1* (Fibronectin) that have previously been shown to facilitate *T. pallidum* attachment to the endothelium^41–43^ (Supplementary Table 1).

We also observed genes encoding ECM modifying proteins to be both up- and downregulated, indicating the dysregulation of endothelial ECM homeostasis during *T. pallidum* exposure. These genes include those belonging to the primary ECM modifying protein families such as MMP (Matrix Metalloproteinase), *ADAM* (A Disintegrin And Metalloproteinases), *ADAMTS* (A Disintegrin And Metalloproteinases with ThromboSpondin Motifs), and *ADAMTSL* (ADAMTS-Like proteinases). (Supplementary Table 1). We also observed dysregulation of the plasmin regulation, including upregulation of the endothelial plasminogen activator *PLAT* (Tissue-type plasminogen activator [also known as *tPA*]). Other related genes included downregulation of the inhibitor of *tPA SERPINB2* (Plasminogen activator inhibitor-2 [also known as PAI-2]), and upregulation of *SERPINE1* (plasminogen activator inhibitor-1 [also known as PAI-1]) (Supplementary Table 1). Notably, *SERPINE1* (PAI-1) expression is induced by TGFβ and Rho/ROCK activation^44^. Conversely, *SERPINB2* is downregulated in response to TGFβ^45^. These findings demonstrate that *T. pallidum* exposure induces significant alterations in expression of key ECM components, regulators of ECM deposition and degradation, and activators/inhibitors of the blood plasminogen/plasmin system. Further, these data highlight the importance of the ECM as a signaling and regulatory interface during *T. pallidum*-host interactions.

### *Treponema pallidum* exposure dysregulated cytoskeletal and junction regulatory pathways and altered HBMEC cellular morphology

In our dataset we observed that pathways involving Rho GTPase activity, as well as those that interact with or modulate Rho signaling, such as RTK, ECM, and integrin signaling, were significantly overrepresented at one or more timepoints (Figure 4, Figure 6D; Supplementary Tables 6-9, 13). Consistent with the overrepresented Rho signaling pathways, Rho GTPases *RHOB*, *RHOD*, *RND1* (Rho family GTPase 1) were upregulated, and *RHOJ* was downregulated (Supplementary Table 1). Further, Rho/Rac/Cdc42 GAPs and guanine nucleotide exchange factors (GEFs) were DE (Supplementary Tables 1). Cdc42 regulators were also DE, including upregulation of the Cdc42-activating protein *CDC42BPG* (Cdc42 binding protein kinase gamma). Several integrins were DE, including upregulation of *ITGΒ3* at 12- and 24-hours, and downregulation of *ITGAX* and *ITGΒ8* (Supplementary Table 1). These findings highlight the dysregulation of Rho GTPases, integrins, and regulators of endothelial junctional dynamics during *T. pallidum* contact.

To further investigate the activation of Rho GTPase signaling in endothelial cells during *T. pallidum* exposure, we performed fluorescent F-actin staining of *T. pallidum*-exposed HBMECs (Figure 7). A subset of HBMECs exposed to *T. pallidum* for 15 and 30 minutes exhibited a contracted cellular morphology, featuring prominent cortical actin rings and slight membrane blebbing (Figure 7A, B). Mean F-actin fluorescence intensity per cell was significantly higher in HBMECs exposed to *T. pallidum* compared to those exposed to basal media. Similarly, HBMECs treated with a biologically relevant concentration of thrombin showed a trend toward increased fluorescence intensity and displayed a contracted morphology with cortical actin ring formation, resembling the response observed in *T. pallidum*-exposed cells (Figure 7A-D). HBMECs exposed to IEC exhibited moderate contraction and a trend toward increased fluorescence intensity, indicating a cellular response to background components in the *T. pallidum* culture media. These findings highlight the pronounced response of HBMECs to viable *T. pallidum.* These alterations had dissipated at the 2-hour timepoint, suggesting a temporal and reversible cellular response to *T. pallidum* engagement (Figure 7C). In summary, our transcriptomic analyses identified integrin/non-integrin, ECM, RTK, and Rho GTPase signaling pathways as significantly overrepresented in *T. pallidum*-exposed HBMECs. Microscopy analyses supported these findings by demonstrating HBMEC F-actin contraction during *T. pallidum* exposure, with viable *T. pallidum* inducing endothelial actin cytoskeletal alterations within minutes of contact.

**Figure 7.**
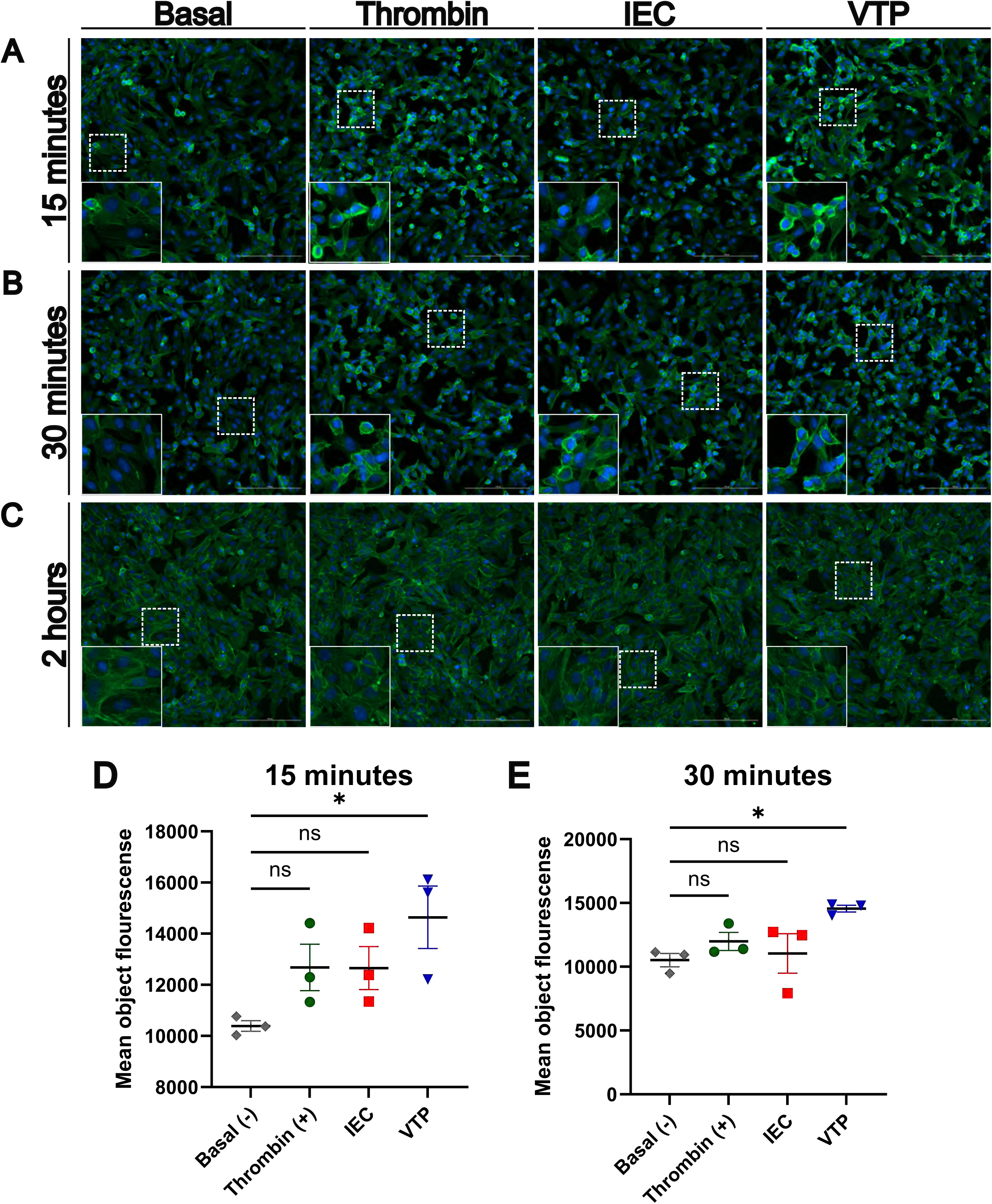
HBMECs exposed to *T. pallidum* displayed a temporal contraction of F-actin, and increased F-actin fluorescence intensity. (A-C) Fluorescent phalloidin F-actin (Green) staining of HBMECs exposed to basal TpCM2 media, or 10 nM bovine thrombin, infection extract control (IEC), and viable *T. pallidum* (VTP). Exposures occurred for (A) 15 minutes, (B) 30 minutes, (C) 2 hours. HBMECs were counterstained with DAPI (Blue). Images were taken in the center of each well using a 20x objective lens. In the lower left side of each panel an inset shows a magnified region of the image, and the location of the original image used for magnification is shown by a dotted box. Scale bar represents 200 μm. (D-E) Plots showing image quantitation; images were taken in an automated 3×3 grid in the center of each well using a 20x objective lens, then stitched linearly to form a single image. Quantified mean object F-actin fluorescence intensity of HBMECs at (D) 15- and (E) 30-minutes of exposure. Each data point represents a biological replicate, which is defined here as an independently treated and imaged well. Significant differences were determined by one-way ANOVA followed by Dunnetts multiple comparison between each condition and the basal media-exposed condition. p-values < 0.05 were considered significant. Error bars indicate the mean with standard deviation. ns = not significant, * indicates a P-value ≤ 0.05.

### *Treponema pallidum* exposure altered interferon, tumor necrosis factor, and inflammatory immune responses

We identified significant dysregulation of immune signaling pathways based on both Sigora and hallmark gene set analyses. Sigora pathway analysis identified the downregulation of chemokine receptor signaling from 4 to 12 hours and the downregulation of IFN gamma and TNF signaling pathways at the 4- and 12-hour timepoints, followed by upregulation of TNF and IFN alpha/beta signaling at 24 hours (Figure 4A-D). Additionally, hallmark gene set analysis identified dysregulation of inflammatory responses from 4- to 24-hours (Figure 6B-D). Downregulated genes within these pathways included monocyte recruiting and activating cytokines/chemokines, such as monocyte chemoattractant protein 1 (*MCP-1* or *CCL2*), *CCL20* (Chemokine C-C motif 20), and colony-stimulating factors (*CSF*)-*1*, *CSF2*, and *CSF3* (Figure 4B-D). IFN immune signaling genes included downregulation of *IRF1* (Interferon regulatory factor 1), and upregulation of *IFNB1* (IFN-β) (Supplementary Table 1).

TNF hallmark gene sets were differentially regulated at all timepoints in the present study, where the “TNF-ɑ signaling through NF-κB” gene set demonstrated both significant up- and downregulation, reflecting the diverse roles of differentially expressed genes within these pathways (Figures 4, 6; Supplementary Tables 6-13). Indeed, genes encoding TNF receptors (TNFRs) were downregulated from 4- to 12-hours and were both up- and downregulated at 24-hours (Supplementary Table 1). TNFR adaptors such as the TRAF (TNF-receptor-associated factor) family exhibited differential expression, primarily at the 24-hour timepoint (Supplementary Table 1). Immune signaling genes downstream of both IFN and TNF, such as the NF-κB factors *TCIM* (Transcriptional and Immune Response Regulator), *NFKB1*, *NFKB2,* and *RelB,* were upregulated, while the NF-κB inhibitor *IKBKE* (Inhibitor of nuclear factor Kappa-B Kinase subunit Epsilon) was downregulated (Supplementary Table 1). Collectively these findings indicate *T. pallidum* exposure alters TNF, IFN signaling, NF-κB signaling and the endothelial immune response to *T. pallidum*.

### Endothelial cell death and viability signaling is altered during *T. pallidum* exposure

In this study, we also found cell death signaling pathways to be overrepresented in both Sigora and hallmark gene set analyses. The apoptosis hallmark gene set was downregulated at 4- and 12-hours and was simultaneously up- and downregulated at 24-hours, though upregulation was more significant (Figure 6B-D). Many pro-apoptotic factors within these pathways were DE, including upregulation of pro-apoptotic genes *PMAIP1* (Phorbol-12-Myristate-13-Acetate-Induced Protein 1 [also known as Noxa]), *BBC3* (Bcl-2 Binding Component 3 [also known as Puma]), and *BCL2L11* (Bcl-2 like 11 [also known as *BIM*]), and downregulation of the anti-apoptotic factor *BCL2A1* (Supplementary Table 1). Sigora pathway analysis identified overrepresentation of pathways related to cell death signaling through p53 at the 24-hour timepoint, where DE genes involved in p53-mediated cell death include upregulation of pro-apoptotic genes *TP73* (Tumor Protein 73), and *TP53INP1* (Tumor Protein p53 Inducible Nuclear Protein 1). Finally, key apoptosis activators *CASP8* (Caspase 8) and *CASP1* were downregulated at 24-hours. These findings demonstrate the dysregulation of apoptotic signaling through intrinsic cell death pathways, such as p53, BH3-only, and through interference with CASP8-regulated cell death signaling.

## Discussion

In this study we investigated the molecular landscape of *T. pallidum*-host interactions through global time-course transcriptomic analyses of HBMECs exposed to viable *T. pallidum.* A prominent transcriptional signature that was observed in this study was endothelial to mesenchymal transition (EndMT), a cellular transformation whereby endothelial cells transition across, or between, intermediary mesenchymal phenotypes^14,46^. Overrepresentation of EndMT pathways or hallmark gene sets, or factors that induce this response, were observed at all timepoints in the study. Further, hallmark gene sets of cellular processes involved in EndMT were overrepresented at the 24-hour timepoint, namely angiogenesis and myogenesis. Consistent with these findings, endothelial markers were downregulated, while mesenchymal markers were upregulated in HBMECs exposed to *T. pallidum*^14,47^. TF enrichment analysis, which identifies TFs that may be responsible for the observed changes in gene expression, indicated that highly connected networks of EndMT-related TFs were enriched across all timepoints. These networks include the essential EndMT-driving TF Snail^21,23^, which was increased in expression on both the transcript and protein levels in HBMECs exposed to *T. pallidum*. Snail upregulation has been shown to promote endothelial permeability and barrier traversal of other bloodborne pathogens^18,19^, and the processes of EndMT and EMT have been proposed to be critical for disrupting endothelial barrier integrity during infectious disease-induced inflammatory conditions^47,48^. In a non-infectious disease context, EndMT promotes tumor cell metastasis during, and prior to, tumor cell-endothelial engagement, facilitating invasion of tumor cells into tissues^49^. Shared mechanisms of dissemination between the metastasis of cancer cells and invasive bacteria have been proposed previously^42,50,51^, and it is plausible that *T. pallidum* employs convergent mechanisms to disseminate.

Fibrosis and endothelial fibrotic dysregulation, as observed in obliterative endarteritis^52^ or syphilis-induced vasculitis (Heubner’s arteritis)^53,54^, are common manifestations of syphilis that can lead to vascular occlusion, tissue ischemia, and infarction^52^. Fibrotic dysregulation of endothelial sites^55,56^ and retinal vasculitis/fibrosis^57,58^ are frequently reported during meningovascular syphilis. Endothelial phenotype modifications have also been documented, where corneal endothelial cells develop a fibroblast-like phenotype with an altered ECM composition^59^. EndMT is an important contributor to vascular and fibrotic disease^14,60^, including though dysregulation of the ECM. The identification of an overarching EndMT transcriptional profile in this study supports clinical observations of endothelial fibrotic involvement during syphilis and identifies a potential functional role for EndMT in weakening cell-cell junctions, *T. pallidum* dissemination, and syphilis disease manifestations.

Pathway overrepresentation analysis revealed significant alterations in EndMT-inducing signaling pathways and hallmark gene sets, including canonical and non-canonical TGFβ, SMAD, β-catenin, NOTCH, NF-κB, and RTK pathways. Indicators of TGFβ activation that induce or potentiate EndMT^14,35,61^ were upregulated, including *NOTCH1*, *IL-11*, *POSTN*, and *EDN1*. Also upregulated were *Nectin-4*, an adherens junction component that promotes EMT through β-catenin/Wnt^62,63^, and *TCIM* that positively regulates β-catenin, endothelial permeability, and MAPK immune signaling^64^. TGFβ negative-feedback regulation, which is activated downstream of TGFβ pathway activation^31^, was evident through elevated expression of inhibitory *SMAD7*, SMAD negative regulators *PMEPA1*, *ID1*, *ID2*, and *ID3*, and decreased expression of TGFβ activators *SMAD3, TGFBR2,* and *TGFBR3*. In parallel, upregulation of *PDGFB*, *PGF*, and *VEGFA* indicated that growth factor activity is altered during *T. pallidum*-endothelial engagement.

These findings are consistent with previous investigations showing that individual *T. pallidum* proteins induce EndMT-related signaling pathways within host cells. Responses include Tp1038 (TpF1) eliciting VEGF/growth-factor-like activity on human umbilical vein endothelial cells (HUVECs)^65^, and Tp0136 inducing fibronectin-mediated integrin-β1 signaling and subsequent microvascular endothelial cell migration^66^. Immune secretion studies have also shown that HBMECs exposed to *T. pallidum* demonstrate increased VEGF secretion^13^. These findings contextualize the observation that rabbits immunized with a tri-antigen vaccine cocktail comprised of vascular adhesin Tp0751^67^ and conserved regions of select *T. pallidum* repeat (Tpr) proteins^68^ exhibit decreased TGFβ transcripts in primary chancres and reduced *T. pallidum* dissemination following *T. pallidum* challenge, compared to challenged unimmunized rabbits^69^. Reduced TGFβ expression would dampen EndMT-promoting signaling at primary chancre sites, thereby reducing the pathology associated with inflammation and endothelial remodeling and attenuating *T. pallidum* dissemination in immunized animals.

Genes and pathways that regulate ECM organization were significantly enriched in the current study, including genes encoding the MMP, ADAM, and ADAML family of matrix proteinases. Previous investigations reported that *T. pallidum* influences the MMP/Tissue inhibitors of metalloproteinases (TIMP) equilibrium in differentiated THP-1 cells^70^, and that *T. pallidum* proteins Tp0136 and Tp0574 (Tp47) alter the MMP/TIMP balance in human dermal vascular smooth muscle cells (HDVSMCs)^71^ and HUVECs^11^. Notably, Tp0136 was shown to alter MMP expression through the PI3K and MAPK signaling cascades^71^, pathways involved in non-canonical TGFβ signaling^14^ that were also found to be altered in the present study. These findings also support the previous report that HBMECs exposed to *T. pallidum* displayed reduced abundance of multiple ECM proteins^13^.

Reports in the literature demonstrate that integrin, ECM signaling, and Rho GTPase regulatory pathways can be exploited by invasive pathogenic bacteria to promote epithelial or endothelial barrier traversal^72^, whereby pathogenic bacteria utilize ECM components as bridging molecules to engage host receptors^73,74^. Growth factor and ECM signaling are cooperative, since ECM constituents can coordinate integrin and RTK signaling^75^. In the current study, integrin interactions, ECM organization, and Rho GTPase pathways were overrepresented in HBMECs exposed to *T. pallidum*. The ability of *T. pallidum* and its constituent proteins to bind ECM components has been well-established^10,41–43,67,76^. Previous work demonstrated that *T. pallidum* fibronectin binding proteins can signal through integrins^66^, and that *T. pallidum* adherence to endothelial cells and fibronectin is reduced in the presence of peptides containing the arginine-glycine-aspartic acid (RGD) cell binding motif found in fibronectin^77^. Integrins were DE in this study, including upregulation of RGD motif-binding integrin *ITGB3*, which is involved in host-pathogen signal transduction^78,79^. Relatedly, we observed that *T. pallidum*-exposed HBMECs displayed increased F-actin signal intensity and a rounded morphology alongside a prominent cortical actin ring, which are hallmarks of RhoA activation^80^. Activation of RhoA downstream of *T. pallidum*-ECM interactions and subsequent enhancement of endothelial traversal has been previously proposed^13,81^, and it has been shown that select *T. pallidum* proteins induce F-actin reorganization through Rho-associated protein kinase (ROCK)-regulated signaling in HUVECs^82^. In a similar study, human dermal lymphatic endothelial cells (HDLECs) exposed to the pathogenic spirochete *Leptospira interrogans* sv. Copenhageni displayed increased F-actin localization and intensity around the cell periphery^83^.

Although *RhoA* was not DE in this study, multiple Rho family members were DE, including *RhoB* and *RND1* as the most significantly upregulated. RhoB functions independently to promote endothelial permeability by decreasing cell-cell contacts, and together with RhoA induces endothelial permeability downstream of endothelial activation^84,85^. Increased RhoB expression also decreases VE-Cadherin localization and accumulation at cell junctions^84,85^. RND1, which is induced by VEGF and TGFβ^86,87^, also disrupts adherens junctions and promotes endothelial cell rounding^88^. These observations provide mechanistic insight into the prior observation that *T. pallidum* modifies endothelial VE-cadherin architecture^9^, and highlights the modulatory effect that *T. pallidum* has on endothelial cytoskeletal and junctional signaling pathways.

The delayed-type hypersensitivity (DTH) response, which is dependent on activation of macrophages, is important for clearing local *T. pallidum* infection^52,89^. In the current study, we detected significant downregulation of *MCP-1* (*CCL2*), *CSF1*, *CSF2*, and *CSF3*. These observations corroborate previous findings where *T. pallidum*-exposed HBMECs displayed reduced secretion of MCP-1 and reduced protein expression of CSF1^13^, and where activated THP-1 macrophages displayed reduced MCP-1 secretion following exposure to *T. pallidum*-derived antimicrobial peptides^90^. MCP-1 and CSF proteins are critical for inducing the DTH response, as these cytokines mediate monocyte recruitment from the bloodstream to sites of endothelial inflammation^91,92^, with subsequent maturation of monocytes to activated macrophages within tissue sites. Due to the importance of DTH responses in clearing local *T. pallidum* infection^52,89^, reduced expression of these cytokines during *T. pallidum* contact might dampen monocyte recruitment and macrophage activity, aiding in *T. pallidum* immune avoidance and persistence. Additionally, IFN immune responses were highly overrepresented and downregulated in the dataset. Given the importance of IFN signaling in anti-viral immunity, the observed downregulation of IFN signaling upon *T. pallidum* engagement with HBMECs could provide additional context for the frequent occurrence of HIV-*T. pallidum* co-infections^93^.

Symptoms stemming from endothelial, mucosal, and tissue destruction are observed at all stages of syphilis and have features characteristic of the process of necroptotic cell death^3,52^. In the current study, apoptosis and TP53-regulated and apoptotic cell death signaling pathways were overrepresented, while necroptosis signaling pathways were not, despite a previous proteomic analysis detecting necroptosis overrepresentation when *in vivo T. pallidum* was exposed to HBMECs^13^. However, necroptosis regulatory factors were DE, including downregulation of the key necroptosis regulator *CASP8* which directs apoptotic signaling to necroptosis when inhibited^94^. It is possible that the *in vitro T. pallidum*-endothelial cell system used in the current study lacks the full complement of factors required to induce a complete necroptotic response within endothelial cells exposed to *T. pallidum*, and confirmation of this prediction must await further studies.

Our investigations herein expand understanding of the endothelial cellular responses induced upon exposure to *T. pallidum*. However, there are limitations to our experimental approach. This study focused on the brain microvasculature, and endothelial cells of different anatomical origin may respond differently to *T. pallidum*. Pericytes, astrocytes, and other cell types contribute to the formation and function of the endothelial and blood-brain barriers, and since the current study investigated HBMECs in monoculture, it may not be representative of the holistic response of endothelial cells to *T. pallidum in vivo.* Further, delineating whether the endothelial responses observed are the result of a protective response raised by the host against *T. pallidum* infection, or a *T. pallidum*-induced manipulation of the host response to enhance pathogenesis of the bacterium, is not easily determined. Finally, the IEC media used in this study is an imperfect control, as it is more inflammatory than the background basal media^13^. This may account for lack of statistical significance in microscopy analyses in F-actin fluorescence intensity between the IEC and VTP treated groups, since inflammatory responses and residual *T. pallidum* components may affect endothelial cell behaviour. However, due to the complex nature of the *T. pallidum* culture systems^95,96^, this control provides the best comparator to measure cellular responses raised specifically to viable *T. pallidum*.

This study provides novel molecular insights into the global host endothelial response to *T. pallidum* and highlights the importance of the ECM for *T. pallidum* pathogenesis. Due to the importance of EMT and EndMT in fetal development and fibrotic disease^14,47^, this study identifies the potential role of EndMT in syphilis disease manifestations observed in infectious and congenital syphilis. Indeed, our findings may further the understanding of HIV-syphilis co-infection since current and previous^13^ observations show that *T. pallidum* exposure downregulates IFN responses integral to both anti-viral immunity and macrophage-mediated clearance of *T. pallidum*. Future multi-omic and data integration investigations focused on the disease stage-specific host response to *T. pallidum,* specifically investigating systemic changes in the host transcriptome, proteome, and metabolome, will further enhance understanding of *T. pallidum*-host interactions and may provide insight into syphilis vaccine development.

## Acknowledgements

We would like to acknowledge Jenna Fleetwood for their assistance with this project. We would also like to acknowledge Travis Blimkie for their assistance with the GEO database. Additionally, we want to acknowledge Drs. Diane Edmondson and Steven Norris for their contribution to the field of syphilis research through the development of an *in vitro* culture system.

## Author contributions

SW: Conceptualization, Formal analysis, Writing – original draft, Writing – review & editing, Data curation, Investigation, Methodology. MG: Formal analysis, Writing – review & editing. AG: Investigation, Writing – review & editing, Methodology. AR: Investigation, Writing – review & editing, Methodology. KL: Writing – review & editing, Conceptualization. RF: Methodology. RH: Conceptualization, Methodology, Writing – review & editing. AL: Conceptualization, Formal analysis, Writing – review & editing. CC: Conceptualization, Formal analysis, Writing – review & editing, Funding acquisition, Project administration, Supervision, Writing – original draft.

## Resource availability

### Lead contact

Further information and requests for resources and reagents should be directed to and will be fulfilled by the lead contact, Caroline Cameron

### Materials availability

This study did not generate new or unique reagents.

### Data and code availability

All sequencing data are archived on GEO (accession: GSE281329) and is publicly available as of the date of publication.

This paper does not report original code.

Any additional information required to reanalyze the data reported in this paper is available from the lead contact upon request.

## Methods

### Treponema pallidum growth

Outbred male specific pathogen-free (SPF) New Zealand White rabbits (3.0-3.5 kg, Charles River Laboratories, Ontario, Canada) with nonreactive syphilis serological tests (VDRL and FTA-ABS) were used for *in vivo* propagation of *T. pallidum* subsp. *pallidum* Nichols strain as previously described^96^. All rabbits were fed antibiotic-free food and water, and were housed at 18-20°C. Animal studies were approved by the local institutional review board under protocol 2020-024 and were conducted in strict accordance with standard accepted principles as set forth by the Canadian Council on Animal Care (CCAC), National Institutes of Health, and the United States Department of Agriculture in facilities accredited by the American Association for the Accreditation of Laboratory Animal Care and the CCAC. Institutional biosafety approval was obtained under biosafety certificate 13170-010. *Treponema pallidum* was cultured *in vitro* with Sf1Ep cells as previously described^95^ in a HeraCell Vios 160i incubator (Thermo Fisher Scientific, San Jose, CA), with the modification that treponemes were dissociated from Sf1Ep cells using trypsin-free dissociation media^97^ for 30 minutes at 34°C in 1.5% O_2_, 5% CO_2_, 93.5% N_2_ to maintain the integrity of the *T. pallidum* outer membrane^95^. Bacteria were quantitated by darkfield microscopy (Nikon Eclipse E600; Nikon Canada, Mississauga, Ontario) using a Petroff-Hauser counting chamber (Hauser Scientific, Horsham, PA). *In vivo* grown *T. pallidum* was used for RNA-seq assays, and *in vitro* grown *T. pallidum* were used for western blot and microscopy assays.

### Viable *T. pallidum* (VTP) and Infection Extract Control (IEC) sample preparations

*In vivo Treponema pallidum* subsp. *pallidum* was extracted in *Treponema pallidum* culture medium 2 (TpCM2) formulated as previously described^95^, or harvested *in vitro* as described above. Viable *T. pallidum* (VTP) and infection extract control (IEC) samples were prepared as described previously^13^. Briefly, extracts were centrifuged to remove rabbit testicular material and diluted equivalent to working *T. pallidum* concentrations. The viable treponeme suspension contained in one supernatant (designated viable *T. pallidum* [VTP]) was kept for endothelial co-incubation analyses as described below. The second supernatant was further processed to remove *T. pallidum* through sterile filtration with a 0.22 μm cellulose acetate syringe filter (Avantor, Allentown, PA) to create an optimal comparator control (designated infection extract control [IEC]) that contained a level of rabbit testicular or Sf1Ep protein background mirroring the VTP sample. Removal of *T. pallidum* from the IEC sample was confirmed by darkfield microscopy (Nikon Eclipse E600; Nikon Canada, Mississauga, Ontario) and *T. pallidum*-specific qPCR, which measured a 49x reduction in *T. pallidum* FlaA copies in the VTP versus IEC samples.

### Endothelial culture and endothelial-*T. pallidum* exposure conditions

Human cerebral brain microvascular endothelial cells (hCMEC/d3; Cedarlane, Burlington, ON), also referred to as HBMECs, were grown to 90% confluence in 6-well tissue culture plates (Corning, Corning, NY), at 5% CO_2_ in EndoGRO-MV complete culture medium (Millipore, Etobicoke, ON) at 37°C in 5% CO_2_ in a Forma Series II incubator (Thermo Fisher Scientific). Each well was observed via an Olympus CKX41 inverted microscope for successful HBMEC growth prior to incubation at 34°C in a microaerophilic environment of 1.5% O_2_, 5% CO_2_, 93.5% N_2_ with 3 mL of either *in vivo* grown VTP (3.0 x10^7^ *T. pallidum* per well), or equivalent dilutions of IEC for 45 minutes, 4, 12, and 24 hours. At each timepoint, there were 5 replicates of VTP- and 5 replicates of IEC-exposed HBMEC sample wells. Therefore, individual wells are defined as separate biological replicates. At each timepoint prior to cell lysis and RNA harvesting, *T. pallidum* organisms were visually observed for motility and viability via darkfield microscopy (Nikon Eclipse E600).

Following treponemal viability assessment, HBMEC cells were quickly washed 3× in cold PBS and incubated in a 1:1 ratio of Qiagen RNAprotect Cell and Qiagen RNAprotect Bacteria (Qiagen, Toronto, ON) for 10 minutes on ice. Cells were pelleted at 10,000 x *g* for 10 minutes, resuspended in 1 mL of RNAprotect cell, and RNA was extracted using the Qiagen RNeasy kit according to manufacturer instructions, with the addition of an on-column DNA digestion using RNase-Free DNase (Qiagen). To prevent RNA degradation, 1 µL of SUPERase-In RNase inhibitor was added to the eluted RNA (Qiagen).

### RNA integrity analysis, cDNA generation, RNA sequencing, and data analysis parameters

Prior to cDNA library generation, RNA integrity of each sample was assessed by an Agilent Bioanalyzer 2100 on an RNA 6000 nanochip (Agilent; Santa Clara, CA), where all RNA samples exceeded an RNA integrity value of 8. Samples then underwent PolyA enrichment using NEBNext Poly(A) mRNA Magnetic Isolation Module (catalog no.: E7409L, NEB; Ipswich, MA). Strand-specific cDNA library preparation was generated at the same time for each sample from polyA-purified RNA with a Roche KAPA HyperPrep Kit (Roche; Basel, Switzerland), followed by the addition of unique 8bp NGS RNA adaptors to identify samples during multiplexed sequencing (NEXTFLEX; PerkinElmer, Woodbridge, ON), generating cDNA with unique tags for each individual sample. cDNA libraries were amplified using adapter primers, followed by purification beads according to manufacturer instructions (NEXTFLEX; PerkinElmer). Amplified DNA quality was assessed on a bioanalyzer using a high sensitivity DNA chip (Agilent) to assess the fragmentation profile and confirm the absence of adaptor-specific primer dimers. cDNA library concentrations were assessed using Qubit QuantIT HS DNA kit (Thermo Fisher Scientific). Each sample was normalized to 4 nM and pooled. Pooled samples underwent paired-end sequencing with a read length of 150bp on an Illumina HiSeqX (Illumina; San Diego, CA, USA) at the Michael Smith Genome Sciences Center (BC, Canada). Sequence quality was assessed using FastQC v0.12.1 and MultiQC v1.13^98^. The FASTQ sequence reads were aligned to the hg19 human genome (Ensembl GRCh38.98) using STAR v2.7.10b^99^ and mapped to Ensembl GRCh38 transcripts. Read-counts were generated using htseq-count (HTSeq 2.0.2)^100^. All data processing and subsequent differential gene expression analyses were performed using R version 4.4.0 and DESeq2^101^ version 1.14.1. Genes with very low counts (with less than 10 counts) were pre-filtered and removed *in silico*. Differentially expressed genes were identified using paired analysis with the Wald statistics test and filtering for any genes that showed ± 1.5-fold-change (FC) with adjusted *p*-values ≤ 0.05 (cut-off at 5% false discovery rate) as the threshold. Pathway overrepresentation analyses were completed using the Reactome database^25^ annotations and Sigora (v3.1.1)^24^ pathway overrepresentation, and the molecular signature database^26^ (MSigDB) hallmark gene set analysis was performed using PathlinkR^102^. The packages maSigPro (v1.74.0)^103^, and the ClusterProfiler R package (v4.10.0)^104^ were used for figure generation. For Sigora and MSigDB analyses, q-values were set at 0.05 using the Benjamini-Hochberg (BH) correction, and pathways or cellular compartments with a corrected p-value ≤ 0.05 were considered significant. Transcription factor analysis was completed using the Chea3 integrated mean ranking algorithm^20^ for each timepoint.

### Western Blotting

Confluent HBMECs in 6-well plates were exposed to *in vitro* grown VTP (3×10^7^ *T. pallidum* per well), equally diluted IEC, or basal TpCM2 media for 12 and 24 hours with 3 biological replicates (defined as individual wells) per treatment condition. Cells were lysed for 30 minutes on ice with gentle agitation in RIPA lysis buffer containing EDTA-free Protease inhibitor cocktail set 3 (Calbiochem, San Diego, CA) and PhosSTOP phosphatase inhibitor (Roche, Mississauga, ON) according to the manufacturer’s instructions. For western blotting, protein concentrations were determined using a BCA assay (Thermo Fisher Scientific). Whole cell lysate (13.5 µg per lane) was subjected to SDS-PAGE using Bolt 12% acrylamide gels (Thermo Fisher Scientific) and transferred to PVDF membrane (Millipore) via wet-transfer at 400 mA for 2 hours, and blocked with Intercept TBS blocking buffer (Licor, Lincoln, NE). Rabbit Anti-Snail (CD15D3; 1:1000; Cell Signalling Technology, Danvers, MA; 3879S) and mouse anti-GAPDH (1:1000; Abcam, Ab8245) were used as primary antibodies, while the secondary antibodies used were goat anti-rabbit IgG IRDye 800CW (1:20,000) and goat anti-mouse IgG IR Dye 680RD (1:20,000; Licor). Detection and analysis were completed on a Licor Odyssey CLx using Licor Image Studio version 5.2. Data was tested for normality using a Shapiro-Wilks tests which determined that the data was consistent with normal distribution, and statistical analysis to determine increases in Snail abundance was completed using One-way ANOVA followed by Tukey’s multiple comparisons testing.

### *Treponema pallidum*-endothelial co-incubation and F-actin staining

hCMEC/d3 endothelial cells were seeded into black wall μClear cell culture-treated bottom 96-well culture plates (Greiner Bio-one, Monroe, NC; 655090) at 24,000 cells/well and grown overnight as described above to achieve 90-95% confluence. *In vitro* cultivated *T. pallidum* was cultured as described above, then VTP and IEC samples were generated as described above and diluted to 4.34×10^6^ cells/mL in TpCM2 media, or equivalent dilution for IEC. For *T. pallidum*-HBMEC co-incubation, HBMEC media was removed, then replaced with 100 μL of VTP, IEC, basal TpCM2 media, or 10 nM bovine thrombin in TpCM2 (Thermo Fisher Scientific; RP-43104) in triplicate per condition, resulting in a multiplicity of infection (MOI) of 30 for VTP samples or equivalent dilution for IEC. Incubations were completed for 15 minutes, 30 minutes, or 2 hours after the addition of *T. pallidum,* and all incubations were performed at 34°C in a microaerophilic environment of 1.5% O_2_, 5% CO_2_, 93.5% N_2_ in a HERAcell vios 160i incubator (Thermo Fisher Scientific). At the specified timepoints, *T. pallidum-*containing supernatants were removed and assessed for motility via darkfield microscopy, where *T. pallidum* motility remained greater than 80% for all timepoints. Simultaneously, HBMECs were washed 2x in pre-warmed PBS, fixed in 3.7% formaldehyde in PBS without methanol for 10 minutes at RT, and washed 3× in pre-warmed PBS. Cells were permeabilized in 0.1% Triton 100-X for 5 minutes, washed 3× in pre-warmed PBS, and blocked in PBS 1% bovine serum albumin (Thermo Fisher Scientific) for 1 hour at 37°C. Alexa fluor 488 Phalloidin (Thermo Fisher Scientific; A12379) stocks were resuspended in DMSO, then diluted to a 1x working solution in PBS with 3μM DAPI counterstain (4′,6-diamidino-2-phenylindole; Sigma Aldrich). Fifty microliters of staining solution was added to each well, the plates were incubated at RT for 20 min, washed 3× in warm PBS, and 100 μL PBS was added to each well for imaging.

### Cell imaging and image processing

All cell imaging was performed on a Cytation 5 Imaging Reader (Agilent) with a 20x objective. Representative images are shown in Figure 7. For statistical analyses wells were imaged in the centre of each well in a 3×3 field of view square. Images underwent image pre-processing and image deconvolution, after which they were stitched into a single image using linear blending in Gen5 (Agilent; v3.12). All imaging was automated and completed at the same time, and with identical imaging conditions. HBMECs were identified (defined as “objects”) and were counted using the DAPI filter as a primary mask (7-35μm). To determine HBMEC cellular boundaries, a secondary mask was set by expanding the primary DAPI mask (110 μm) in the GFP (FITC; F-actin) channel using the threshold in mask method (FITC; F-actin); touching objects were split, objects contacting the border of the image were excluded, and gaps in masks were filled. Using the secondary mask, F-actin fluorescence intensity was determined within the boundaries of each object, the fluorescence intensity was calculated for each defined object, and the average object fluorescence intensity was calculated for each well. Significant increases in fluorescence intensity were determined by comparing mean object fluorescence intensity from 3 biological replicates (defined as individual wells) per sample condition. Data was tested for normality using a Shapiro-Wilks tests which determined that the data was consistent with normal distribution, and significance was determined using One-way ANOVA followed by Dunnett’s multiple comparison, using HBMECs exposed to basal media as the baseline for statistical comparison.

## Supplemental information titles

**Supplementary Table 1.** List of DE gene fold changes and p-values from 45-minutes to 24-hours. Official gene symbol and ENSEMBL gene IDs are shown. Differentially expressed genes were identified using paired analysis with the Wald statistics test and filtering for any genes with a fold-change cutoff of ±1.5 and adjusted p-value < 0.05 (cut-off at 5% false discovery rate) as the threshold for significance.

**Supplementary Table 2.** Chea3 transcription factor enrichment analysis of the DE genes at the 45-minute timepoint. Transcription factors were ranked using the integrated mean ranking algorithm. Overlapping genes are genes identified to be targeted by the respective transcription factor.

**Supplementary Table 3.** Chea3 transcription factor enrichment analysis of the DE genes at the 4-hour timepoint. Transcription factors were ranked using the integrated mean ranking algorithm. Overlapping genes are genes identified to be targeted by the respective transcription factor.

**Supplementary Table 4.** Chea3 transcription factor enrichment analysis of the DE genes at the 12-hour timepoint. Transcription factors were ranked using the integrated mean ranking algorithm. Overlapping genes are genes identified to be targeted by the respective transcription factor.

**Supplementary Table 5.** Chea3 transcription factor enrichment analysis of the DE genes at the 24-hour timepoint. Transcription factors were ranked using the integrated mean ranking algorithm. Overlapping genes are genes identified to be targeted by the respective transcription factor.

**Supplementary Table 6.** Sigora pathway overrepresentation analysis of up- and downregulated DE genes at the 45-minute timepoint. Pathways with a Bonferroni-corrected p-value < 0.05 were considered significant. The q-value for Bonferroni correction was 0.05.

**Supplementary Table 7.** Sigora pathway overrepresentation analysis of up- and downregulated DE genes at the 4-hour timepoint. Pathways with a Bonferroni-corrected p-value < 0.05 were considered significant. The q-value for Bonferroni correction was 0.05.

**Supplementary Table 8.** Sigora pathway overrepresentation analysis of up- and downregulated DE genes at the 12-hour timepoint. Pathways with a Bonferroni-corrected p-value < 0.05 were considered significant. The q-value for Bonferroni correction was 0.05.

**Supplementary Table 9.** Sigora pathway overrepresentation analysis of up- and downregulated DE genes at the 24-hour timepoint. Pathways with a Bonferroni-corrected p-value < 0.05 were considered significant. The q-value for Bonferroni correction was 0.05.

**Supplementary Table 10.** Molecular signature database hallmark gene set overrepresentation analysis of DE genes at the 45-minute timepoint. Pathways with an adjusted p-value < 0.05 were considered significant. Hallmark gene sets may be concurrently up- and down-regulated.

**Supplementary Table 11.** Molecular signature database hallmark gene set overrepresentation analysis of DE genes at the 4-hour timepoint. Pathways with an adjusted p-value < 0.05 were considered significant. Hallmark gene sets may be concurrently up- and down-regulated.

**Supplementary Table 12.** Molecular signature database hallmark gene set overrepresentation analysis of DE genes at the 12-hour timepoint. Pathways with an adjusted p-value < 0.05 were considered significant. Hallmark gene sets may be concurrently up- and down-regulated.

**Supplementary Table 13.** Molecular signature database hallmark gene set overrepresentation analysis of DE genes at the 24-hour timepoint. Pathways with an adjusted p-value < 0.05 were considered significant. Hallmark gene sets may be concurrently up- and down-regulated.

